# *Seqpare*: a self-consistent metric of similarity between genomic interval sets

**DOI:** 10.1101/2020.04.05.026732

**Authors:** Selena C. Feng, Nathan C. Sheffield, Jianglin Feng

## Abstract

**Summary:** Searching genomic interval sets produced by sequencing methods has been widely and routinely performed; however, existing metrics for quantifying similarities among interval sets are inconsistent. Here we introduce *Seqpare*, a self-consistent and effective metric of similarity and tool for comparing sequences based on their interval sets. With this metric, the similarity of two interval sets is quantified by a single index, the ratio of their effective overlap over the union: an index of *zero* indicates unrelated interval sets, and an index of *one* means that the interval sets are identical. Analysis and tests confirm the effectiveness and self-consistency of the *Seqpare* metric.

**Availability:** https://github.com/deepstanding/seqpare

## INTRODUCTION

Functional genomic data are often summarized as interval sets and deposited in public repositories (e.g., UCSC, ENCODE, Roadmap, GEO, SRA etc.). Identifying relationships among sequences and searching through widely available sequence data are routine tasks in genomic research. A fundamental operation in genomic/epigenomic analysis is comparing two interval sets, and many algorithms and tools have been developed for this purpose (Alekseyenko & Lee, 2007; Kent, 2002; Quinlan & Hall 2010; Li, 2011; Neph etal, 2012; Cormen et al. 2001; Jalili et al., 2019; Feng et al., 2019). These methods are based on computing the total number of intersections (overlaps) between the two interval sets. To compare a query interval set with multiple interval sets in a genomic sequence database, searching tools LOLA (Sheffield & Bock, 2016) and GIGGLE (Layer et al., 2018) calculate two values — Fisher’s exact *p*-value and the odds-ratio based on the total number of intersections — and use them as the similarity score to rank the search results. These similarity metrics have proven useful for determining relationships among interval sets, but also have some flaws. First, calculating the Fisher’s exact test results requires building a contingency table, but determining its values is not straightforward. The *p*-value and odds-ratio for two intervals sets (with number of intervals *N*_1_ and *N*_2_) are calculated from four numbers, namely, the number of intersections between the two sets, *n*, the number of intervals in set 1 that do not overlap an interval in set 2, *N*_1_ - *n*, the number of intervals in set 2 that do not overlap an interval in set 1, *N*_2_ - *n*, and the number of intervals that are not present in either set, *m*. Determining the value of the fourth number *m* is not straightforward; in LOLA, it depends on the definition of a “universe set” that is not objectively defined, whereas GIGGLE estimates *m* from the two interval sets. Second, the total number of overlaps *n* does not necessarily reflect similarity since intervals can have very different lengths (often in the range of 1~10^5^ base pairs) and two very different intervals may intersect by only a few base pairs. This can result in inconsistency of the metrics: a comparison between two identical interval sets may have a larger *p*-value or smaller odds-ratio than a comparison between two different interval sets (see example cases and analysis in the next section). More strikingly, since one interval may contain or cover other intervals in an interval set, depending on how the overlaps are computed, *n* can be larger than *N*_1_ and/or *N*_2_, i.e., *N*_1_-*n* and/or *N*_2_-*n* can be negative, which leads to both the *p*-value and odds-ratio being undefined—another potential source of inconsistency. Third, the Fisher’s exact-based metrics require two values (*p*-value and odds-ratio) but neither is a *direct* measurement of the similarity: *p*-values are sensitive to the total number of regions and can range as low as 10^-200^ for large genomic interval sets, and odds-ratios are sensitive to small numbers; and neither metric directly informs on *how similar* the two sets are. Last, the *p*-value calculation is computationally expensive for genomic interval sets, particularly when the number of intervals is large (up to 10^9^). To overcome these weaknesses of the Fisher’s exact-based metrics, we developed *Seqpare*, a self-consistent metric for quantifying the similarity among genomic interval sets.

## METHOD

### Seqpare metric

The *Seqpare* metric uses a single index to quantify the degree of similarity *S* of two interval sets with number of intervals *N*_1_ and *N*_2_. Similar to the *Jaccard* index, the *Seqpare* metric is directly defined as the ratio of the total effective overlap *O* of the two interval sets over the union *N*_1_+*N*_2_ - *O*:

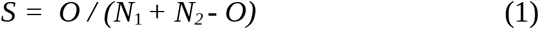

For two intervals *v*_1_ in set 1 and *v*_2_ in set 2, the similarity *s* is defined as:

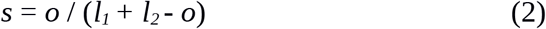

where *o* is the length of the intersection and *l*_1_ and *l*_2_ are the lengths of *v*_1_ and *v*_2_ respectively. Definition 2 is the *Jaccard* index for individual intervals: *o* represents the effective overlap of the two intervals and *s* takes values in the range of [0, 1]: *s* = 0 indicates that there is no overlap between the two intervals, and *s* = 1 means that the union equals the overlap so *v*_1_ and *v*_2_ are identical. Then the total effective overlap *O* for the two interval sets can be calculated by adding up the similarities of all mutual best matching (MBM) pairs:

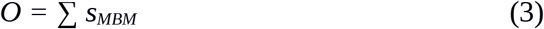

A MBM pair is defined as a pair of intervals *v*_1_ and *v*_2_ that fulfill the following conditions: among all intervals in set 2 that intersect *v*_1_, *v*_2_ matches *v*_1_ the best, i.e., the similarity *s* between *v*_1_ and *v*_2_ is the highest among those intersections; and among all intervals in set 1 that intersect *v*_2_, *v*_1_ matches *v*_2_ the best. Clearly, if two intervals only intersect each other, then they form an MBM pair. In Figure 1a, the two long intervals (the 1st in set 1 and the 1st in set 2) only intersect each other (intersection pair *ip*_1,1_), so they form an MBM pair; similarly the two short intervals *ip*_2,2_ form another MBM pair. For intervals that involve multiple intersections, we define a relatively simple and strict rule to find the MBM pairs: find and choose the first MBM pair that has the highest *s* among all involved intersection pairs, then find and choose the next MBM pair that has the highest *s* from the rest of the intersection pairs (excluding all pairs that involve the intervals that are already chosen), and so on until there are no more intersection pairs. In Figure 1b, there are 3 intersection pairs: *ip*_1,2_ with *s*=1/5, *ip*_1,3_ with s=1/9, and *ip*_2,3_ with s=1/5. So the first MBM pair is either *ip*_1,2_ or *ip*_2,3_ depending on which one is found first. If *ip*_1,2_ is chosen as the first MBM pair, then *ip*_1,3_ will not be considered since interval 1 in set 1 is already chosen, and then we have only *ip*_2,3_ left, which is the second MBM pair. We get the same result if *ip*_2,3_ is chosen as the first MBM pair. In Figure 1d, there are 6 *ip*s and two *MBM pairs: ip*_1,1_ and *ip*_3,2_ both with *s*=1, where interval 2 in set 1 (*i*_2,1_) does not have a match. Note that interval *i*_2,1_ matches best with interval *i*_1,2_, but *i*_1,2_ does not match best with *i*_2,1_, so they are not the *mutual* best matching pair.

**Figure 1.**
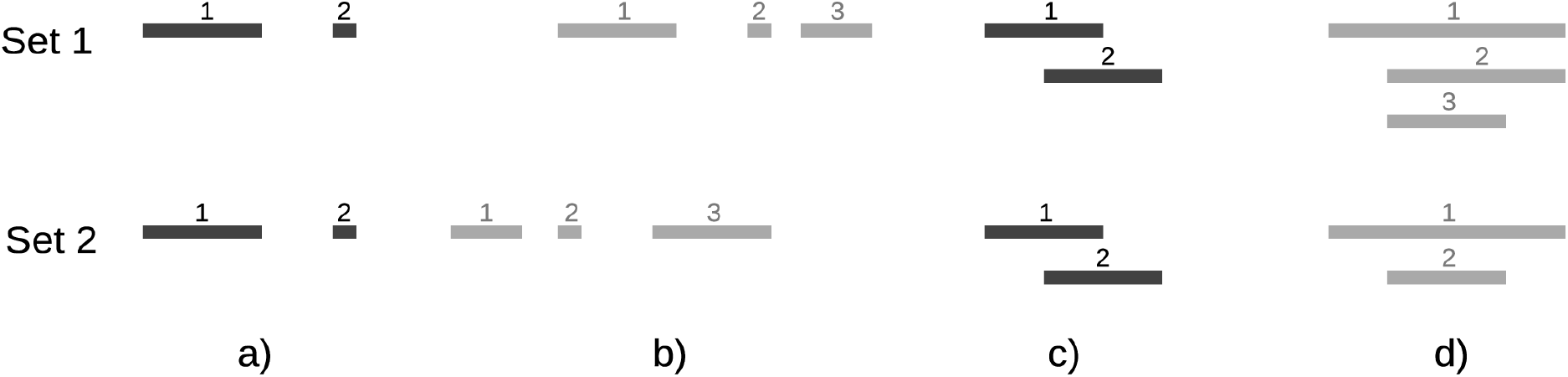
Example cases for illustrating the Seqpare similarity metric. The length ratio of the short interval to the longer intervals are 1: 5 in a), 1: 3: 5 in b), and 2:3:4 in d). The total number of overlaps n is 2 for a), 3 for b) (interval 1 in Set 1 intersects two intervals in Set 2), 4 for c) and 6 for d). The p-value in case b is smaller than that in case a. N_1_ - n and N_2_ - n are both negative in cases c and d.

Since the number of total matching pairs ≤ Min(*N*_1_, *N*_2_) — the minimum of *N*_1_ and *N*_2_ — and *s* is in range of [0, 1], we obtain *O* ≤ Min(*N*_1_, *N*_2_), and *S* takes a value in the range of [0, 1]. If *S* is zero, then there is no matching pair, and vice versa; if *S* = 1, then *N*_1_ = *N*_2_ = *O* (the two sets are equivalent), and vice versa. And, because each *s* is the *mutual* best match, *O* is symmetric (the amount of overlap between set 1 and set 2 is the same as that between set 2 to set 1) and so is *S*.

In Figure 1a, *O_a_*=2 and *S_a_*=1, which is correct because the two sets are identical. In Figure 1b, *O*_b_=2/5 and *S*_b_=1/14, which is expected since the two sets are very different. The Fisher’s exact approach is inconsistent here: the *p*-value in 1*b* is smaller than that in 1*a* although the two sets in 1*b* are very different while those in 1*a* are equivalent. Assuming that the number of intervals *N* in the ‘universe set’ is 100, then Fisher’s exact test contingency table is [(2, 0), (0, 98)] in 1*a* and [(3, 0), (0, 97)] in 1*b*, which gives *p*_a_ = 2.02×10^-4^ and *p*_b_ = 6.18×10^-6^ respectively. The odds-ratio is ∞ in both cases. In cases *c* and *d*, *N*_1_ - *n* and *N*_2_ - *n* are all negative, so it is not conceptually appropriate to use Fisher’s exact test to calculate the *p*-value and odds-ratio.

### Implementation

The implementation of the *Seqpare* metric is simple. The searching for MBM pairs is deterministic and it can be implemented by directly following the description in the above section. The *Seqpare* code is built on top of the AIList (Feng et al., 2019) software written in C.

## RESULT

### A test with real genomic interval sets

To test *Seqpare* and compare it with the Fisher’s exact-based metrics, we took 100 interval sets from a UCSC database and used one interval set, *affyGnf1h*, as a query to search over the database. Because the database contains the query set, *affyGnf1h* should have the highest similarity score. Table 1 shows part of the result. Interval set *affyGnf1h* indeed ranks first with maximum similarity 1 when using *Seqpare*, but it ranks 94^th^ out of 100 when using the *p*-value and ranks last when using the odds-ratio. This happens because *N*_1_-*n* and *N*_2_-*n* are both negative (n=16686, *N*_1_=*N*_2_=12158). Given this inconsistency, GIGGLE sets negative *N*_1_-*n* and *N*_2_-*n* to zero to calculate the *p*-value, and to one to calculate the odds-ratio. The *Seqpare* indices for other interval sets are all small (<0.03) because the average effective overlap of an intersection pair in those sets is about 0.1 or less, i.e., they are very different from the query set affyGnf1h; however, all of the *p*-values are so small (e^-200^), which suggests that the *p*-value is not a meaningful similarity index for these genomic interval sets. This search takes 6m30s for *Seqpare* and 15m32s for GIGGLE. All computations were carried out on a computer with a 2.8GHz CPU, 16GB memory, and an external SSD hard disk. The complete results can be found at the same site as the software.

**Table 1.** Comparison of Seqpare and GIGGLE similarity metrics: partial list of the search results from a collection of 100 interval sets, which contains the query set affyGnf1h.

| *Seqpare* metric |  | *p*-value and odds-ratio |  |  |  | Interval dataset<br>file name |
| --- | --- | --- | --- | --- | --- | --- |
| rank | similarity | rank_p | *p*-value | rank_or | odds-ratio |  |
| 1 | 1.0 | 94 | $1.543 \text{ e}^{-197}$ | 100 | $9.046 \text{ e}^{-16}$ | affyGnf1h |
| 2 | 0.029 | 19 | $8.328 \text{ e}^{-201}$ | 1 | 25.25 | ccdsGene |
| 3 | 0.026 | 35 | $2.743 \text{ e}^{-200}$ | 3 | 4.425 | allenBrainAli |
| 4 | 0.022 | 39 | $3.372 \text{ e}^{-200}$ | 32 | $7.972 \text{ e}^{-11}$ | affyU133Plus2 |
| 5 | 0.021 | 4 | $5.403 \text{ e}^{-202}$ | 2 | 17.24 | affyU133 |

## CONCLUSION

We have shown that the Fisher’s exact test approach may be not the most appropriate test statistic for comparing similarity among interval sets. While the approach has been shown to be successful for many questions, we have demonstrated how it can break down for a variety of reasons, such as very similar interval sets, within-set containment, widely varying interval lengths among sets, or small effective overlaps. In contrast, *Seqpare* is a self-consistent metric for quantifying the similarity of two interval sets that addresses these concerns. *Seqpare* is the first rigorously defined metric for comparing two sequences based on their interval sets. In addition to the metric itself, our *Seqpare* software tool provides functions for both searching and mapping large-scale interval datasets. We anticipate that this approach will contribute to novel results for interval set searching.

